# Target engagement in human motor cortex induced by constant sinusoidal and amplitude-modulated transcranial AC stimulation

**DOI:** 10.1101/2025.09.29.679395

**Authors:** Mitsuaki Takemi, Mads JA Madsen, Janine Kesselheim, Hartwig R Siebner

**Affiliations:** Danish Research Centre for Magnetic Resonance, Department of Radiology and Nuclear Medicine, Copenhagen University Hospital Amager and Hvidovre, Hvidovre, Denmark; Graduate School of Advanced Science and Engineering, Hiroshima University, Hiroshima, Japan; Department of Neurology, Copenhagen University Hospital Bispebjerg and Frederiksberg, Copenhagen, Denmark; Department of Clinical Medicine, Faculty of Health and Medical Sciences, University of Copenhagen, Copenhagen, Denmark

**Keywords:** tACS, TMS, EEG, corticospinal excitability, neuromodulation

## Abstract

Transcranial alternating current stimulation (tACS) is a noninvasive technique for modulating brain oscillations. While sinusoidal tACS (sin-tACS) delivers current at a constant amplitude, amplitude-modulated tACS (AM-tACS) uses a high-frequency carrier modulated by a low-frequency envelope. We systematically compared the acute effects of sin-tACS at theta (5 Hz), alpha (10 Hz), beta (20 Hz), and gamma (140 Hz) frequencies, and AM-tACS (140 Hz carrier frequency modulated at theta, alpha, or beta) on corticospinal excitability.

Healthy participants received 2 mA peak-to-peak tACS to the primary motor hand area (M1_HAND_) via a bipolar montage (M1_HAND_–Pz). Each tACS block consisted of ten 30-second stimulation periods interleaved with 6-second pauses, followed by 11 minutes without stimulation. Corticospinal excitability was assessed during each block using single-pulse transcranial magnetic stimulation (TMS), delivered at the peak and trough of tACS.

MEP amplitudes were generally larger at the trough. Only beta sin-tACS and theta AM-tACS significantly modulated corticospinal excitability. Beta sin-tACS increased MEP amplitudes in early stimulation epochs, while theta AM-tACS facilitated MEPs in later stimulation epochs and this facilitation persisted briefly after stimulation ended. Participants with stronger early responses to beta sin-tACS also tended to show greater delayed effects with theta AM-tACS. These excitability changes during tACS were not predicted by simulated electric field strength. A follow-up EEG experiment revealed that beta sin-tACS increased beta power over left sensorimotor cortex, while theta AM-tACS decreased beta power over midline parietal cortex. These EEG changes were restricted to tACS pauses.

The results show that sin-tACS and AM-tACS can both modulate corticospinal excitability, but functional changes differ in temporal dynamics, frequency specificity, and cortical region engagement.

## Introduction

Transcranial alternating current stimulation (tACS) is a non-invasive brain stimulation technique in which low-intensity current is applied to the brain at alternating polarity with zero offset (Antal et al., 2008). Most tACS applications use a steady sinusoidal frequency (i.e., sin-tACS) inducing an oscillating current in the brain. Depending on the frequency and amplitude, the alternating current can interact with specific ongoing endogenous neural rhythms of the human brain (Ali et al., 2013; Frohlich & McCormick, 2010; Helfrich et al., 2014; Johnson et al., 2020; Krause et al., 2019). These effects are thought to mediate beneficial tACS effects on normal brain function like attention, memory, and executive control or disrupted brain functions in neurological and neuropsychiatric disorders including schizophrenia, attention-deficit/hyperactivity disorder, and Parkinson’s disease (Frohlich et al., 2015; Grover et al., 2023; Riddle & Frohlich, 2021; Wischnewski et al., 2023).

Several studies have combined sin-tACS with single-pulse transcranial magnetic stimulation (TMS) of the primary motor hand area (M1_HAND_) and measured the mean amplitude of motor evoked potentials (MEPs) to assess tACS-related changes in corticospinal excitability. While a seminal study failed to show consistent effects, later studies have shown immediate (during sin-tACS) and outlasting (after the end of sin-tACS) effects on resting corticospinal excitability at various frequencies, including sin-tACS within the beta-band and high gamma-frequency range (Feurra et al., 2011; Guerra et al., 2016; Moliadze et al., 2010; Schutter et al., 2011; Wischnewski et al., 2019). Several studies have also observed phase-dependent differences in MEP amplitudes during tACS administration, suggesting that the sinusoidal current alterations may contribute to cortical fluctuations in excitability (Guerra et al., 2016; Nakazono et al., 2016; Raco et al., 2016; Schilberg et al., 2018).

Amplitude-modulated tACS (AM-tACS) is a newer form of tACS during which the amplitude of a high-frequency carrier wave (e.g., >100 Hz) is modulated with a lower-frequency envelope, typically in the range of natural brain rhythms (e.g., theta: 4–8 Hz, alpha: 8–12 Hz, beta: 13–30 Hz) (Negahbani et al., 2018; Witkowski et al., 2015). This differs from sin-tACS, where the stimulation is applied directly at the frequency of interest (e.g., a 6 Hz sinusoid without any high-frequency carrier) (Barzegar et al., 2025). The mechanisms of AM-tACS are not yet fully understood but are likely to differ from standard sin-tACS. Endogenous brain oscillations often exhibit cross-frequency coupling, where high-frequency rhythms (e.g., beta, gamma) are modulated by the phase or amplitude of slower oscillations (e.g., theta, alpha) (Helfrich & Knight, 2016; Sauseng et al., 2019). One common form of this is phase–amplitude coupling (PAC), in which the amplitude of faster oscillations fluctuates with the phase of a slower oscillation (Daume et al., 2017; Yanagisawa et al., 2012). This coupling is thought to play a fundamental role in cognitive processes such as working memory, attention, sensorimotor integration, and memory consolidation (Canolty et al., 2006; Lisman & Jensen, 2013; Mölle & Born, 2011; Yanagisawa et al., 2012). AM-tACS is well-suited to engage these dynamics, because it reproduces endogenous PAC by embedding a high-frequency carrier (e.g., gamma) within a low-frequency envelope (e.g., theta). This temporal structure may more effectively interact with neural computations relying on such temporal dynamics (Alekseichuk et al., 2016). By contrast, standard sin-tACS targets a single frequency and does not capture this nested organization. Studies that have directly compared the physiological effects of sin-tACS and AM-tACS on the human cortex are scarce. A recent psychophysical study reported higher phosphene thresholds for AM-tACS than for sin-tACS (Hsu et al., 2022). Unlike sin-tACS, the perceived phosphene flicker rate under AM-tACS was not related to the carrier frequency (Hsu et al., 2022).

In this study, we directly compared the physiological effects of conventional sin-tACS and AM-tACS. We applied seven different patterns of stimulation targeting the M1_HAND_ with a bipolar montage. We probed corticospinal excitability with single-pulse TMS of M1_HAND_ during (online) and immediately after (offline) tACS sessions. In a separate experiment, we measured electroencephalography (EEG) to gain insights into tACS-induced alterations in cortical resting activity. Results reveal differences in mechanism of action but also a close relationship in intra-individual responsiveness to both forms of tACS.

## Materials and Methods

### Study design and participants

We conducted two experiments in healthy volunteers to assess the acute effects of seven different tACS protocols targeting the left M1_HAND_. The first experiment employed single-pulse TMS to probe corticospinal excitability during and shortly after each tACS protocol, while the second experiment used EEG to capture short-term effects of each tACS protocol on cortical resting activity. The TMS-tACS and TMS-EEG experiments were separated by more than one week to avoid potential carry-over effects.

Twenty healthy volunteers (6 female; age range: 18–39 years) participated in experiment 1 in which single-pulse TMS and tACS were combined. Two participants declined participation in the subsequent EEG-tACS experiment. All participants reported 6.5–8 hours of sleep the night prior and were right-handed, as determined by the Edinburgh Handedness Inventory with a laterality index ranging from 69 to 100 (Oldfield, 1971). None were regular smokers or had a history of neurological, psychiatric disorders, or contraindications to TMS or MRI. All participants received detailed oral and written information and provided written informed consent. The study was carried out under the Helsinki declaration and approved by the local ethics committee of the capital region of Denmark (H-15017238).

### MRI acquisition

All participants underwent whole-brain structural MRI using a magnetization prepared rapid gradient echo sequence (voxel size: 0.85 mm^3^ isotropic, flip-angle: 8°, TR/TE: 6/2.7 ms), a 3T Achieva scanner and 32-channel receiving head coil (Philips, Best, Netherlands). The individual structural brain scan was used to spatially guide the placement of the tACS electrodes in both experiments and to navigate TMS coil positioning in experiment 1. The brain scans also enabled us to calculate the electric field distribution induced by tACS for each participant.

### Standard and amplitude modulated tACS

In both experiments, conventional sin-tACS was applied at 5, 20, and 140 Hz, and amplitude-modulated tACS (AM-tACS) was delivered using a 140 Hz carrier frequency modulated at 5 and 20 Hz. The first experiment included two additional tACS conditions, namely conventional sin-tACS at 10 Hz and AM-tACS at 140 Hz carrier frequency with an amplitude modulation at 10 Hz (Fig. 1A). Each tACS protocol was administered in separate blocks, presented in a pseudo-randomized, counterbalanced order across participants. A single tACS block consisted of ten 30-second epochs, interleaved by a 6-second pause, resulting in a repeated 30 seconds "on" / 6 seconds "off" pattern. Each tACS block was followed by an 11-minute tACS-free period to assess potential aftereffects and to minimize carry-over effects between subsequent tACS conditions.

**Figure 1.**
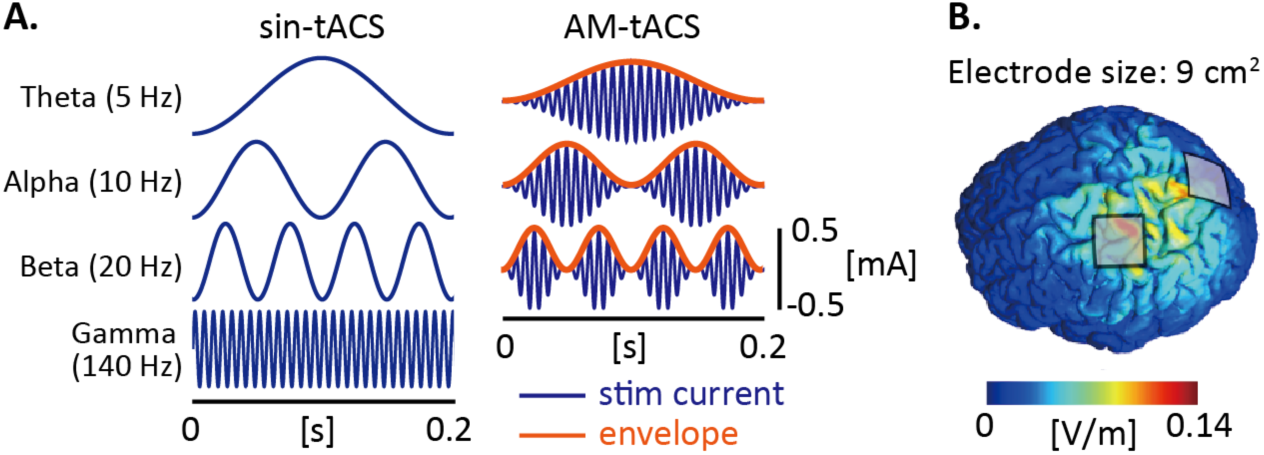
(A) Illustration of the seven stimulating currents tested in the present study, including conventional sinusoidal transcranial alternating current stimulation (sin-tACS) and amplitude-modulated tACS (AM-tACS). (B) Electrode montages and the induced electrical field of a representative participant.

In all tACS conditions, stimulation was delivered at 2 mA peak-to-peak without DC offset. To reduce the impact of peripheral co-stimulation at onset and offset, the current was ramped up and down over 4 seconds. Stimulation was applied via two 9 cm² square silicone-rubber electrodes (NeuroCare Group GmbH, Munich, Germany) placed over the left M1_HAND_ and the Pz position of the international 10–20 EEG system (Fig. 1B). Electrodes were affixed to the scalp using Ten20 conductive paste following skin preparation with NuPrep Skin Prep gel (Weaver and Company, Aurora, CO, USA), and secured with net caps (Spes Medica, Genova, Italy). The outer corners of both electrodes were registered in the neuro-navigation system to enable subsequent electric field modeling and to verify that the M1_HAND_ hotspot was located within the inner electrode boundary. Electrode impedance was kept below 5 kΩ prior to stimulation. The stimulation waveforms were generated using Signal software (version 4.11) and transmitted to the stimulator via a Micro1401-4 analog-to-digital converter (Cambridge Electronic Design, Cambridge, UK).

### Experiment 1: tACS effects on corticospinal excitability

We combined tACS with single-pulse TMS to test for frequency-specific effects of tACS on corticospinal excitability. Conventional sin-tACS was applied at 5, 10, 20, and 140 Hz, and AM-tACS at 140 Hz with sinusoidal amplitude modulation at 5, 10, and 20 Hz, resulting in seven tACS conditions. Single pulse TMS was delivered to left M1_HAND_ during and after each tACS condition to probe immediate (online) and lasting (offline effects) of the various tACS conditions on corticospinal excitability.

A MagPro X100 Option and a MC-B70 figure-of-eight coil were used to stimulate the left M1_HAND_ (MagVenture, Farum, Denmark). The figure-of-eight coil was placed tangentially over the scalp with the handle directed posteriorly in a 45° angle to the sagittal midline, and monophasic pulses inducing a posterior-anterior current in the cortex were administered. Correct TMS coil position over M1_HAND_ was monitored continuously by a frameless stereotaxic neuro-navigation system (Localite, Bonn, Germany), and the root mean square of the difference between the co-registered landmarks was monitored not to exceed 2 mm to ensure adequate accuracy throughout the experiment.

Motor evoked potentials (MEPs) were recorded using surface electromyographic (EMG) electrodes (Ambu Neuroline 710, Ballerup, Denmark) placed on the right first dorsal interosseous muscle using a bipolar belly-to-tendon montage and a ground electrode mounted at the right processus styloideus ulnae. Participants were seated comfortably in a neck-supported reclined chair with their arms bent and relaxed, resting on a pillow placed in their lap. EMG activity was monitored online throughout the measurements, and participants were told to relax if background EMG activity was present. TMS pulses were separated by an inter-stimulus interval of 6±1 seconds during and after the tACS blocks. The timing of TMS and MEP recordings were controlled by Signal software version 4.11 (Cambridge Electronic Design, Cambridge, UK). The signal was amplified (×1,000) and bandpass filtered (5–2,000 Hz) through a DC amplifier (D360, Digitimer, Hertfordshire, UK), digitized at 5 kHz using an analog-to-digital converter (Micro1401-4) (Cambridge Electronic Design, Cambridge, UK) and stored for offline analysis.

The individual M1_HAND_ hotspot was defined as the scalp position where suprathreshold TMS elicited the most consistent and prominent MEPs while the participant was at rest. This location was marked for later monitoring of the coil position using frameless neuro-navigation (Localite, Bonn, Germany). After placement of the tACS electrodes, the resting motor threshold (RMT) was determined using the maximum-likelihood parameter estimation by sequential testing (ML-PEST) method (Awiszus, 2003), defining MEPs ≥50 μV as responses.

Corticospinal excitability was assessed at baseline and during each of the seven tACS blocks, applying single monophasic TMS pulses to the M1_HAND_ hotspot at 120% resting motor threshold intensity (Fig. 2A). Prior to tACS, we recorded 20 MEPs to establish baseline excitability. Each tACS block consisted of ten 30-second stimulation epochs, separated by 5-second intervals without tACS. During each 30-second epoch, five MEPs were recorded. During the tACS stimulation TMS pulses were semi-randomly timed to the peak (0°) and trough (180°) of the sinusoidal waveform to examine phase-specific modulation of excitability (Fig. 2B). In the conventional sin-tACS condition, 25 TMS pulses were delivered at both the peak and trough of the waveform. For AM-tACS, 16 or 17 TMS pulses were applied across three distinct phases: (1) the peak of the carrier wave at the peak of the envelope, (2) the trough of the carrier wave at the peak of the envelope, and (3) the trough of the envelope. We also recorded a single MEP in each 5-second tACS-off interval, adding 10 inter-epoch MEPs per block.

**Figure 2.**
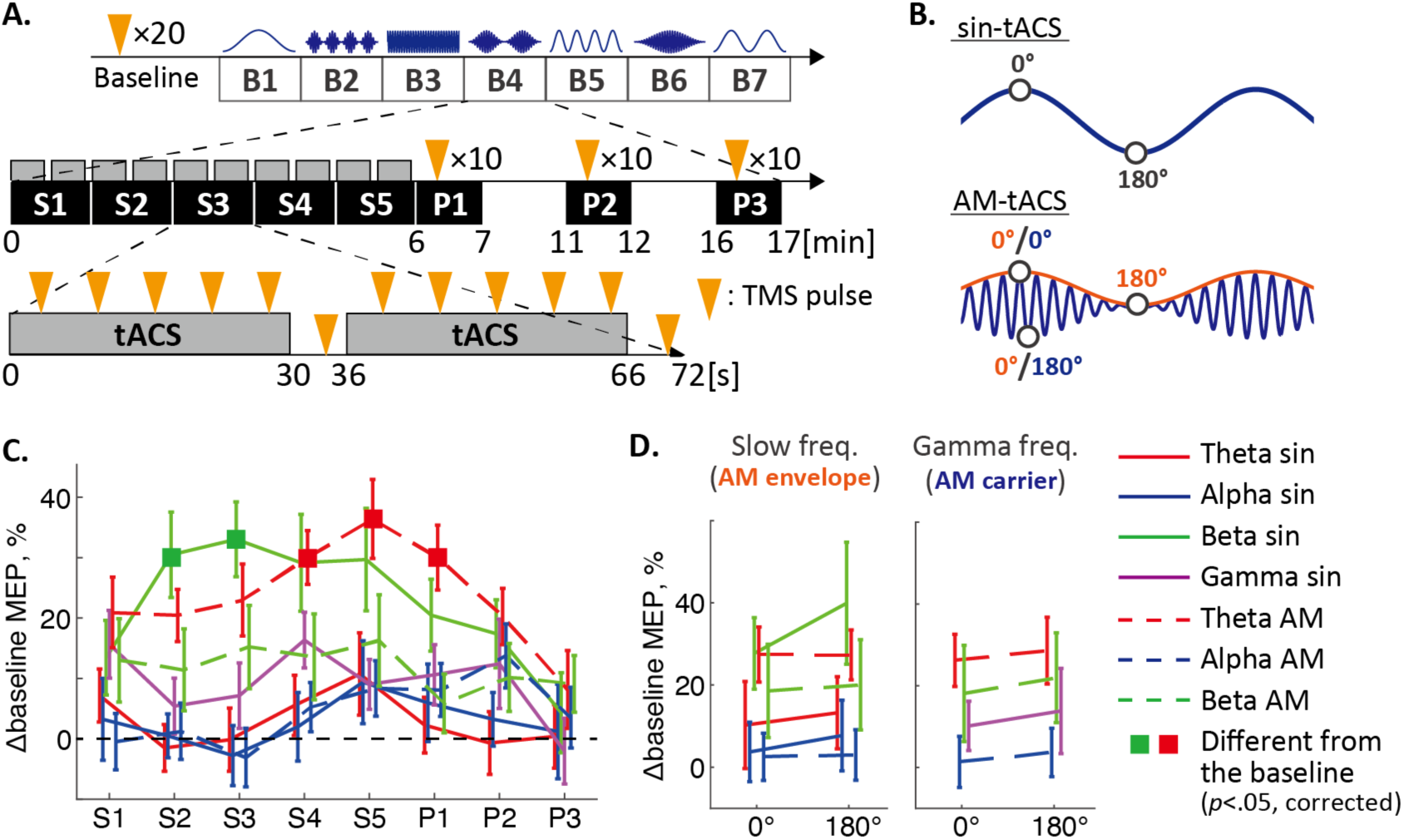
Experiment 1 assessed the effects of the tACS protocols shown in figure 1 on corticospinal excitability. (A) Experimental timeline. The experiment consisting of one baseline block followed by seven tACS blocks (B1−B7), each using different stimulation waveforms. The order of the seven tACS patterns was pseudo-randomized and counter-balanced between participants. Each tACS block started with a 6-minute tACS period consisting of ten 30-second epochs. 6-second pauses were interleaved between consecutive blocks, resulting in a repeated 30 seconds "on" / 6 seconds "off" pattern. tACS was followed by an 11-minute tACS-free period to assess potential aftereffects. A total of 90 motor-evoked potentials (MEPs) were obtained during each block. The timing of single TMS pulses is indicated by the orange triangles. Changes in corticospinal excitability were evaluated both online (S1−S5) and offline (P1−P3). (B) Relative timing of TMS pulses during tACS. TMS was given at two phases in the sin-tACS conditions (i.e., peak and trough of the sine wave) and at three phases in the AM-tACS conditions (i.e., peak and trough of the sine wave at maximum of tACS amplitude modulation and at its minimum). (C, D) Changes in corticospinal excitability during the seven tACS blocks quantified by the mean MEP amplitudes normalized to baseline. Error bars reflect standard errors of the mean. Squares in the plot represent time epochs where the MEP amplitudes were significantly larger than the baseline (*p* < 0.05).

To evaluate after-effects, ten MEPs were recorded at three post-stimulation time points: 0–1 min, 5–6 min, and 10–11 min following each tACS block. During the second minute after stimulation, participants completed a visual analog scale (VAS, range: 0–10) to rate tACS-related side effects, including the strength and discomfort of phosphenes, tickling/prickling sensations, and metallic taste.

### Experiment 2: Frontoparietal resting activity during tACS-free periods

This experiment utilized EEG to evaluate whether tACS modulates intrinsic oscillatory activity in the frontoparietal cortex. Previous work has demonstrated that non-linear stimulation-induced artifacts may persist in EEG signals even after artifact suppression procedures, potentially confounding physiological interpretations (Noury et al., 2016; Noury & Siegel, 2017). To avoid this risk, EEG data were acquired exclusively during stimulation-free intervals both within and following tACS application, thereby avoiding contamination by residual artifacts at the stimulation frequency.

Given that neither sin-tACS (sin-tACS) nor amplitude-modulated tACS (AM-tACS) at alpha frequency demonstrated any measurable effect on corticospinal excitability in prior assessments, these frequency conditions were excluded from experiment 2. Instead, we restricted stimulation to conventional sin-tACS at 5 Hz, 20 Hz, and 140 Hz, as well as AM-tACS employing a 140 Hz carrier modulated at 5 Hz and 20 Hz. Hence, the experiment consisted of five tACS-EEG blocks (Fig. 4A). Stimulation waveforms were generated in MATLAB 2016a (The MathWorks, Natick, MA, USA) and delivered to a neuroConn DC-STIMULATOR PLUS (neuroCare Group GmbH) via a PCIe-6353 digital-to-analog converter (National Instruments, Austin, TX, USA).

**Figure 3.**
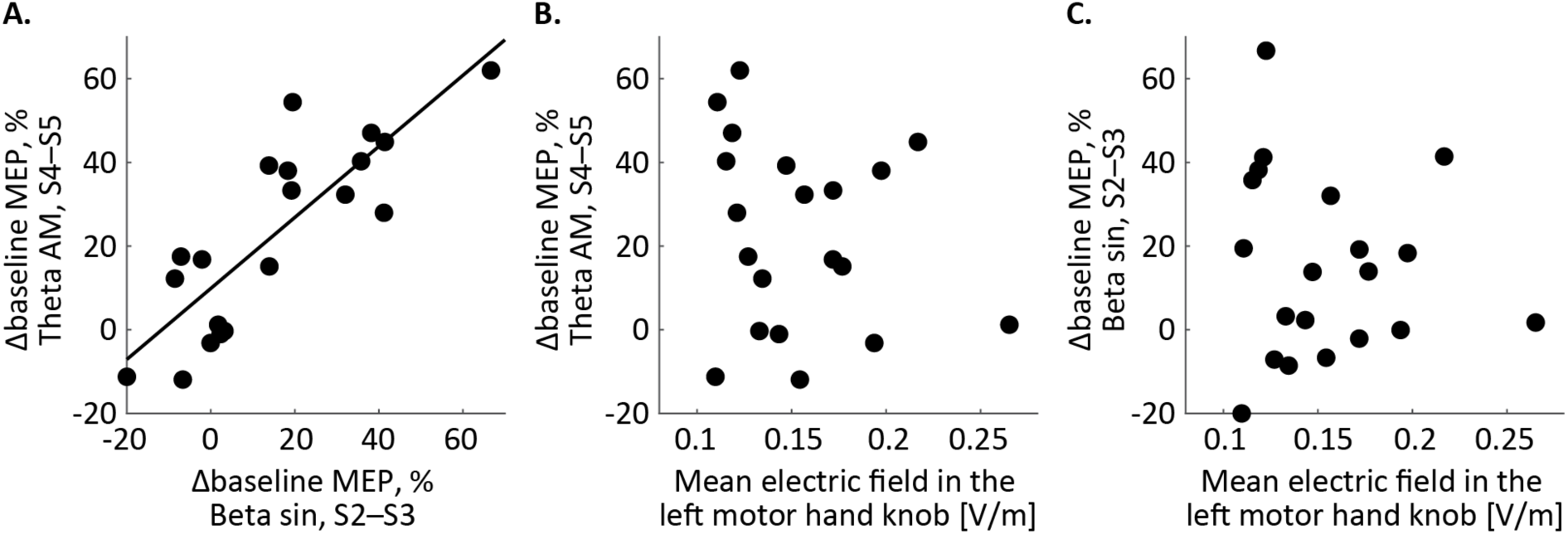
Relationship between changes in MEP and electric field characteristics during different tACS conditions. (A) Scatter plot showing the correlation between the percentage change in MEP amplitude from baseline during the late phase of theta AM-tACS (S4–S5) and the percentage change during the middle phase of beta sin-tACS (S2–S3). Each dot represents an individual participant, and the solid line indicates the line of best fit, demonstrating a positive correlation. (B) Relationship between the mean electric field intensity in the left precentral gyrus, which forms the motor hand knob, and the percentage change in MEP amplitude during the late phase of theta AM-tACS. (C) Relationship between the mean electric field intensity in the left motor hand knob and the percentage change in MEP amplitude during the middle phase of beta sin-tACS.

**Figure 4.**
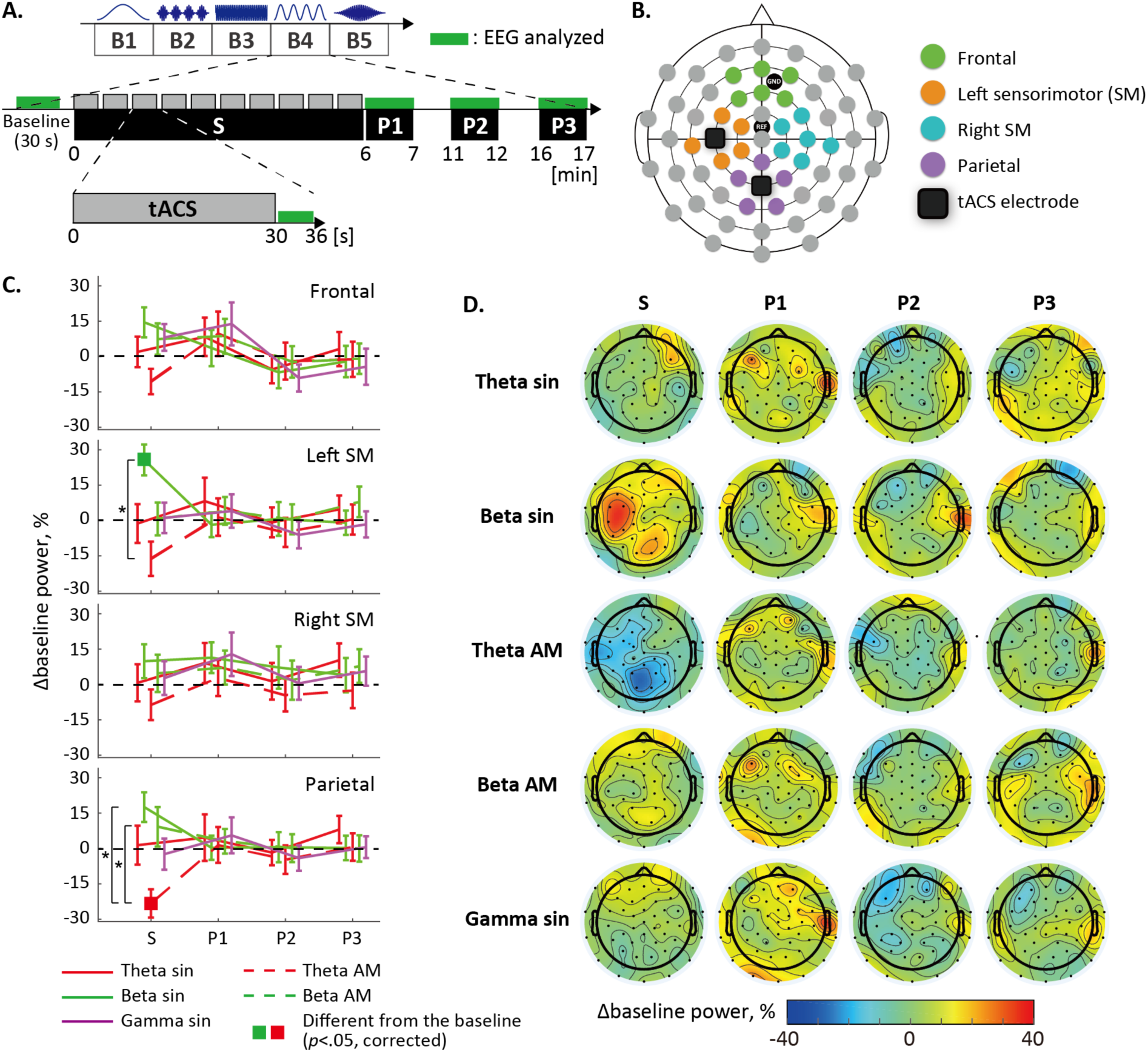
Experiment 2 assessed effects of five different tACS protocols on frontoparietal cortical activity with EEG. (A) Experimental timeline. In this experiment, only five tACS conditions were tested, namely sin-tACS at 5 Hz, 20 Hz, and 140 Hz, as well as AM-tACS employing a 140 Hz carrier modulated at 5 Hz and 20 Hz. Apart from minor adjustment in electrode positioning, the tACS protocol was identical to experiment 1. 30 seconds of baseline EEG measurements were followed by five tACS blocks (B1−B5). The green bars indicate the time epochs during which EEG signals were analyzed. Note that EEG analysis was restricted to periods when tACS was stopped (i.e., in pauses between consecutive tACS epochs and at three epochs after the end of tACS). (B) Positions of EEG and tACS electrodes, with several EEG channels assigned to one of four cortical clusters (frontal, left sensorimotor (SM), right SM, and parietal) for data analysis. (C, D) tACS-dependent changes in beta (19−21 Hz) EEG power relative to baseline. Values shown in Fig. 4C represent the averaged EEG power within the clusters. Error bars reflect standard errors of the mean. Squares in the plot represent time epochs where the EEG power significantly differs from the baseline. *: *p* < 0.05 by post-hoc pairwise comparisons.

Scalp EEG was recorded with a 63-channel equidistant M10 electrode cap (Easycap, Herrsching, Germany) attached to a NeuroOne Tesla system (Mega Electronics, Kuopio, Finland). A total of 63 electrodes, including the cap’s designated ground and reference, were used for data acquisition, excluding the two sites directly beneath the tACS electrodes. The skin was prepared with NuPrep Skin Prep gel (Weaver, Aurora, CO, USA) followed by Abralyt HiCl electrode gel (Easycap, Herrsching, Germany) to ensure optimal conductivity. Electrode impedance was maintained below 5 kΩ throughout the recording session. EEG signals were bandpass filtered between 0.16 and 1,250 Hz, digitized at a sampling rate of 5 kHz, and streamed in real-time to MATLAB 2016a (The MathWorks, Natick, MA, USA). Only EEG data from stimulation-free intervals were retained for offline analysis to avoid contamination from tACS-related artifacts.

To maximize the number of useable EEG channels during concurrent tACS, two square (9 cm²) conductive silicone-rubber stimulation electrodes were positioned beneath EEG electrodes near the M1_HAND_ hotspot and Pz, based on the international 10–20 system (Fig. 4B). While these tACS electrode placements differed slightly from those used in the primary experiment, they still adequately covered the targeted cortical areas. The tACS electrodes were affixed to the scalp using Ten20 conductive adhesive paste (Weaver and Company, Aurora, CO, USA) and stabilized with the EEG cap. Electrode impedance was confirmed to be below 5 kΩ prior to stimulation.

Each tACS-EEG block began with a 30-second baseline EEG recording, followed by ten cycles of 30 seconds of tACS, each interleaved with a 6-second stimulation-free interval. During tACS blocks, EEG data were collected during the central 5 seconds of each stimulation-free period, excluding the initial and final 0.5 seconds to minimize transient effects. In addition to intra-block recordings, EEG was also recorded for one minute at three post-stimulation intervals: 0–1, 5–6, and 10–11 minutes following each tACS block. The sequence of five tACS conditions was pseudo-randomized and counterbalanced across participants to mitigate order effects. Blocks were separated by a minimum one-minute rest period to reduce fatigue.

### Data analysis

#### Motor evoked potentials (MEPs)

EMG data were processed using a custom-made MATLAB script (The MathWorks, Natick, MA, USA). All EMG traces were visually inspected and discarded if any visible background muscle contraction was detected. Peak-to-peak amplitudes of the MEPs for the remaining trials were calculated, and all MEP amplitudes were normalized to the average of baseline MEP amplitudes to assess the relative changes in corticospinal excitability induced by tACS.

To examine the time course of MEP changes, a single tACS block was separated into five online epochs, referred to as S1-S5, and three offline epochs, referred to as P1–3 (Fig. 2A). The mean values of normalized MEP amplitudes were then calculated for each epoch and analyzed using an ANOVA consisting of the two within-subject factors *time epoch* (S1–5 and P1–3) and *tACS condition*. Post-hoc pairwise comparisons were performed using the Bonferroni procedure to correct for multiple comparisons. Bootstrap tests were also performed to examine whether a pattern of tACS significantly modulated corticospinal excitability relative to baseline. At each time epoch of each tACS condition, the normalized MEP values were randomly resampled to create 5,000 different datasets, which were used to calculate 5,000 bootstrapped means. A histogram of the bootstrapped means was then used to test significance with Bonferroni correction for multiple comparisons.

Since the TMS pulse was either given at the peak or trough of the alternating current, we also contrasted MEP amplitudes obtained at the peak or trough of the theta, alpha, beta, and gamma frequencies produced during tACS. For sin-tACS conditions, we calculated mean values of normalized MEP amplitudes separately for the peak and trough of the tACS phases (Fig. 2B). For gamma tACS conditions in which the amplitude of the carrier frequency was modulated, we calculated mean values of normalized MEP amplitudes at the gamma peak and trough, when tACS amplitude peaked as well as at the time point when tACS amplitude was zero (Fig. 2B). The mean MEP values were then tested using an ANOVA with three within-subject factors *phase of tACS* (peak vs. trough), *tACS frequency* (theta, alpha, or beta), and *type of tACS* (sinusoidal vs. amplitude modulated).

To assess modulation of corticospinal excitability at the gamma frequency, we calculated mean values of normalized MEP amplitudes separately for the two tACS phases during gamma sin-tACS and separately for the two carrier wave phases during AM-tACS. The mean MEP values were tested using an ANOVA with two within-subject factors: phase (peak / trough) and envelope frequencies (none / theta / alpha / beta). For all the statistical analysis, the type I error was set to 0.05. JASP 0.18.1 was used for ANOVA and post-hoc pairwise comparisons. Bootstrap tests were performed using MATLAB 2021b (The MathWorks, Natick, MA, USA).

#### EEG data

EEG data analysis focused on EEG activity in the beta and theta band as experiment 1 had only shown changes in corticospinal excitability during sin-tACS at beta frequency and AM-tACS with amplitude modulation at theta frequency.

EEG signals were bandpass filtered between 1–45 Hz using a 4th-order Butterworth filter and down-sampled to 200 Hz. Data were visually inspected to identify and exclude noisy channels, after which the remaining channels were re-referenced using a common average reference. Independent component analysis (ICA) was applied to remove both physiological (e.g., eye blinks, eye movements) and non-physiological artifacts. EEG data were segmented into 1-second epochs and converted to time-frequency representations via FFT using a non-overlapping Hanning window. Power spectra were normalized per channel and frequency to the baseline average of the corresponding tACS block.

To assess frequency-specific EEG modulation, EEG recordings during each block were divided into four epochs, one online epoch (S), corresponding to the pooled epochs S1-S5, and three offline epochs after the end of each tACS block (P1–P3). Normalized EEG power within each epoch was winsorized at the 90th percentile to mitigate outlier effects, then averaged across two defined frequency bands (theta: 4–6 Hz; beta: 19–21 Hz) and four cortical regions, corresponding to groups of EEG channels overlying frontal, parietal, left and right sensorimotor cortex (Fig. 4B). Mean normalized EEG power was analyzed using a repeated-measures ANOVA with within-subject factors: time epoch, cortical region, and tACS condition. Greenhouse-Geisser correction was applied when sphericity was violated, and Bonferroni-adjusted post hoc tests were used for pairwise comparisons. To test for deviations from baseline, bootstrap analyses were performed. For each epoch and tACS condition, normalized power values were resampled 5,000 times to compute bootstrapped means, and significance was assessed using Bonferroni-corrected thresholds derived from the resulting distributions.

### Electric field estimation

Individual electric field strengths were estimated using SimNIBS software v4.1.0 (Thielscher et al., 2015) based on individualized head models generated from T1-weighted MRI scans via the ‘charm’ pipeline. The tACS electrodes were recorded via neuronavigation and imported using custom MATLAB scripts. Electrodes were modeled as 4 mm thick rubber pads (conductivity: 29.4 S/m) with a 2 mm layer of conductive paste (conductivity: 1 S/m). We examined the relationship between electric field strength and tACS-induced changes in corticospinal excitability, as indexed by the MEP amplitudes of the FDI muscle. To this end, we defined the left motor hand knob as the region of interest (ROI), encompassing M1_HAND_ and the adjacent dorsal premotor cortex (Dubbioso et al., 2021).

## Results

### Experiment 1: tACS effects on corticospinal excitability with TMS

#### Frequency-dependent modulation of corticospinal excitability

Figure 2C depicts changes in mean MEP amplitude relative to baseline across tACS conditions. Certain tACS protocols produced transient increases in corticospinal excitability, but effects varied among conditions. A two-factor repeated-measures ANOVA showed a significant interaction between time epoch and tACS condition (F(11.3, 213.8) = 2.23, p = 0.013), showing that the modulation of corticospinal excitability was contingent on both the waveform and the time of MEP measurement relative to tACS delivery.

Post-hoc bootstrap analyses identified two tACS waveforms that significantly increased MEP amplitudes at specific time points. Beta sin-tACS led to a significant increase in MEP amplitude during the early stimulation epochs (S2 and S3; p < 0.05). Conversely, amplitude-modulated tACS with a theta-frequency envelope produced a delayed excitatory effect, with significant MEP increases observed in the later stimulation epochs and persisting immediately post-stimulation (S4, S5, and P1; p < 0.05). No other tACS conditions elicited statistically significant changes in MEP amplitude relative to baseline.

Although the changes in corticospinal excitability during sin-tACS at beta-frequency and AM-tACS at theta-frequency were expressed at different time points during tACS, we explored whether the tACS-induced increases in MEP amplitudes showed a linear relationship across individuals (Fig. 3A). This exploratory analysis revealed a strong positive correlation between the percentage change in MEP amplitude during the late period of theta AM-tACS (S4–S5) and the percentage change during the middle period of beta sin-tACS (S2–S3) (r(18) = 0.831, p < 0.001). This finding suggests that individuals who exhibited stronger corticospinal excitability enhancement during beta sin-tACS also tended to exhibit greater excitability increases during theta AM-tACS.

Importantly, no differences in mean MEP amplitude were detected between the initial (S1) and last epoch (P3) of the same block across all seven tACS conditions. This finding indicates that single tACS protocols did not produce measurable offline aftereffects extending beyond 10 minutes, irrespective of the stimulation waveform.

#### Online tACS effects on corticospinal excitability at tACS peak and trough

We examined whether the phase of the slow components (theta, alpha, beta) of tACS waveform modulated corticospinal excitability by comparing MEP amplitudes elicited at the peak versus trough of the sinusoid (for sin-tACS) or the envelope (for AM-tACS). As shown in Figure 2D left, mean MEP amplitudes were larger when TMS was applied at the trough of the waveform. This peak-trough difference was supported by a three-way repeated-measures ANOVA with within-subject factors of tACS phase, frequency, and type, revealing a main effect of phase (F(1, 19) = 5.04, p = 0.037). Exploratory peak-trough comparisons for each individual tACS condition revealed no significant difference in MEP amplitudes for any tACS conditions, indicating that the main effect of phase reflected a small, consistent trend across conditions rather than being driven by a single frequency or waveform.

Additionally, the ANOVA revealed an interaction between the type of tACS (sinusoidal vs. amplitude modulated) and tACS frequency (*F*(1.75, 33.2) = 8.60, *p* < 0.001), indicating that the effects of tACS on MEP amplitude are determined by both amplitude modulation and frequency. This interaction is consistent with our earlier findings that only theta AM-tACS and beta sin-tACS transiently modulated corticospinal excitability during stimulation. No significant interaction was observed between tACS type and phase (F(1, 19) = 3.16, p = 0.092), nor was there a significant three-way interaction among phase, frequency, and tACS type (F(1.39, 26.3) = 0.406, p = 0.60).

To further evaluate phase-specific effects at higher frequencies, we conducted a separate two-way ANOVA on MEP amplitudes during gamma sin-tACS and during AM-tACS with theta, alpha, beta envelopes (Fig. 2D right). This analysis revealed neither a significant main effect of phase (F(1, 19) = 1.31, p = 0.27) nor a significant interaction (F(1.46, 27.8) = 0.017, p = 0.96). These results suggest that while corticospinal excitability appears to be modulated in a phase-dependent manner, this may be limited to frequencies at or below 20 Hz.

#### Relationship between corticospinal excitability and induced electric field strength

We explored the relationship between electric field intensity in the left motor hand knob and the observed increases in corticospinal excitability modulation for theta AM-tACS and beta sin-tACS (Fig. 3B). The changes in MEP amplitude during theta AM-tACS did not correlate with the mean electric field strength (r(18) = -0.12, p = 0.42). Likewise, beta sin-tACS showed no correlation between MEP changes and electric field intensity in the left motor hand knob (r(18) = -0.11, p = 0.66; Fig. 3C).

#### Sensory co-stimulation effects during tACS

A tickling sensation during at least one tACS condition was reported by 16 participants (Suppl. Fig. 1). A one-way repeated-measures ANOVA with Greenhouse-Geisser correction for non-sphericity revealed no significant main effect of stimulation waveform on the intensity of tickling sensation (F(2.7, 40.3) = 2.02, p = 0.133). Other sensory co-stimulation effects were generally mild and infrequent. Phosphenes were reported by three participants: one during theta-frequency sin-tACS, two during beta-frequency sin-tACS, and one during alpha-frequency AM-tACS. Four participants reported experiencing a metallic taste, distributed across conditions as follows: theta sin-tACS (n = 1), gamma sin-tACS (n = 1), theta AM-tACS (n = 2), and alpha AM-tACS (n = 1). Taken together, these findings indicate no systematic differences in sensory co-stimulation effects across the various tACS conditions tested.

### Experiment 2: Frontoparietal resting activity during tACS-free periods

#### Changes in beta-band EEG power

Relative changes in beta-band EEG power across four cortical regions are depicted in Figure 4C. The effects of tACS on cortical beta oscillations varied as a function of stimulation waveform, cortical region, and time of assessment. A three-way repeated-measures ANOVA with normalized beta power as the dependent variable revealed a significant interaction among tACS condition, cortical region, and time epoch (F(36, 612) = 1.48, p = 0.039).

Post-hoc bootstrap analyses indicated that the two stimulation conditions associated with changes in corticospinal excitability, namely beta sin-tACS and theta AM-tACS, produced distinct and region-specific effects on beta EEG power. Specifically, beta sin-tACS led to a significant online increase in beta power over the left sensorimotor cortex, close to the stimulation electrode (p < 0.05 at S). In contrast, theta AM-tACS resulted in a significant online decrease in beta power over the midline parietal cortex, where the second electrode was positioned (p < 0.05 at S). Topographical analysis of beta power changes between stimulation periods further suggested that these effects may have propagated toward the contralateral electrode site, indicating potential inter-electrode spread of neuromodulatory influence (Fig. 4D). These changes in beta power were transient. They emerged only during tACS application (i.e., in the short pauses interleaved between tACS epochs) and dissipated immediately after stimulation ceased. No significant changes in beta power were observed over the frontal cortex or the contralateral (unstimulated) sensorimotor area, indicating spatial specificity of the observed effects.

#### Changes in theta-band EEG power

Relative changes in theta-band EEG power across four cortical regions are summarized in Supplementary Figure 2. A three-way repeated-measures ANOVA with normalized theta power as the dependent variable revealed a significant main effect of time epoch (F(2.37, 40.3) = 12.3, p < 0.001), indicating temporal variation in theta activity relative to baseline. Changes in theta power were transient, emerging after cessation of tACS. Post-hoc analysis comparisons showed that theta power was significantly higher during P1 compared to P2 and P3 (p < 0.05). However, no significant main effect or interaction involving the factor tACS waveform was observed. This suggests that a transient boost of theta-band power was independent of stimulation pattern and EEG recording site, reflecting a spatially non-specific effect that was shared by the five tACS conditions.

## Discussion

We found that both sin-tACS and AM-tACS exhibit frequency-specific “sweet spots” for increasing corticospinal excitability. With a bipolar pericentral–parietal montage, beta sin-tACS enhanced MEP amplitudes, whereas alpha and theta sin-tACS did not. For AM-tACS with a 140 Hz carrier, excitability increased only when the amplitude was modulated at theta frequency but not at alpha or beta frequency. We first discuss the effects of constant-amplitude versus amplitude-modulated AC stimulation on corticomotor excitability and contrast these schemes and their physiological effects. We will then comment on the lack of consistent aftereffects and the non-specific but weak phase dependency of tACS effects. Finally, we will highlight methodological limitations and identify open questions that warrant further investigation.

### Online effects during tACS administration

The increase in corticospinal excitability during sin-tACS at beta frequency is consistent with previous studies (Feurra et al., 2011; Wischnewski et al., 2019; Pozdniakov et al., 2021). This may reflect alignment of the extrinsically imposed “oscillations” with intrinsic beta oscillations of motor areas that are prevailing in the “resting” motor cortex (Khanna & Carmena, 2015).

Such frequency-specific alignment has been suggested to promote interactions between externally applied and endogenous neural rhythms (Herrmann et al., 2013), which could in turn enhance neuronal excitability during tACS. Supporting this notion, beta sin-tACS was accompanied by increased beta power over the sensorimotor cortex during interleaved pauses. In contrast, AM-tACS with beta-frequency modulation neither enhanced MEP amplitudes nor EEG beta power, indicating that sin-tACS and AM-tACS at beta frequency are not functionally equivalent and likely engage distinct mechanisms.

AM-tACS enhanced corticospinal excitability only when amplitude was modulated at theta frequency. While no prior studies have combined a gamma carrier with low-frequency modulation to target corticomotor excitability, Akkad et al. (2021) applied a related approach by nesting gamma bursts within theta phases. Using a similar montage, they showed that theta–gamma tACS over right M1_HAND_ facilitated motor skill learning with effects persisting for at least an hour. These findings highlight theta–gamma stimulation as a promising strategy for probing and modulating motor system function.

Unlike theta AM-tACS, continuous theta sin-tACS did not reliably increase corticospinal excitability, underscoring mechanistic differences between stimulation modes. Prior work indicates that sin-tACS effects on corticospinal excitability depend on both frequency and cortical state: beta sin-tACS was most effective at rest, whereas theta sin-tACS enhanced excitability during motor imagery (Feurra et al., 2013). This raises the possibility that theta AM-tACS and theta sin-tACS may converge in efficacy under active states such as motor imagery, when the cortex is more susceptible to theta-targeted stimulation.

Responses to tACS varied considerably across participants. Even protocols effective at the group level failed to enhance MEPs in some individuals. This variability aligns with prior evidence of heterogeneous responsiveness to transcranial brain stimulation (Corp et al., 2020; Kasten et al., 2019; Miniussi & Bortoletto, 2025; Ridding & Ziemann, 2010; Ziemann & Siebner, 2015). Despite this variability, we found a strong positive correlation between individual responses to beta sin-tACS and theta AM-tACS. Individuals in whom MEP amplitude increased markedly during one protocol also showed a clear MEP facilitation during the other tACS protocol, whereas weak responders showed little modulation under both tACS conditions. These findings suggest a trait-like cortical susceptibility to tACS. Therefore, future work should not only try to define optimal stimulation parameters at the group level (i.e., sweet spots) but also identify physiological factors that determine overall sensitivity to tACS at these sweet spots.

Both beta sin-tACS and theta AM-tACS enhanced corticospinal excitability during AC stimulation and tACS-induced increases in MEP amplitude were closely correlated at the subject level. Yet, these communalities do not indicate that both tACS protocols increase corticomotor excitability via identical mechanisms. On the contrary, our results point to different mechanisms of actions. First, MEP facilitation showed distinct temporal dynamics. Beta sin-tACS induced an early transient increase in corticospinal excitability, observable within the first 2–3 min of stimulation which spontaneously tapered off despite continuing sin-tACS. This crescendo-decrescendo pattern suggests that beta sin-tACS may concurrently trigger reactive mechanisms over relatively short timescales (minutes), such as compensatory downscaling of synaptic strength or upscaling of inhibition which bring corticospinal excitability back to baseline (Fricke et al., 2011; Li et al., 2014; Misonou, 2010; Romer et al., 2016).

In contrast, theta AM-tACS elicited a delayed enhancement, that first emerged during the later phase of stimulation (3–5 min) and outlasted the end of AM-tACS for a short period. This suggest a more gradual build-up of excitability changes which may be due to the more intermittent periodic pattern of stimulation.

In an additional experiment, we analyzed EEG during short pauses following tACS with minimal artifact contamination. Beta sin-tACS increased beta power in electrodes overlying the stimulated left sensorimotor cortex, compatible with local entrainment. In contrast, theta AM-tACS did not enhance local theta activity but reduced beta power over midline parietal sites, indicating a modulation at a network level beyond the stimulation site. Given the role of theta oscillations in large-scale communication (Solomon et al., 2017), theta AM-tACS may influence corticospinal excitability via frontoparietal connectivity rather than local entrainment, consistent with effects of theta–gamma tACS on motor learning and cognitive control (Akkad et al., 2021; Riddle et al., 2021). Together, these results point to distinct, potentially degenerate mechanisms underlying MEP facilitation with beta sin-tACS versus theta AM-tACS, underscoring the need for further physiological dissection of the basic mechanisms that are engaged by tACS.

### Lack of aftereffects on cortical excitability after tACS administration

Notably, none of the tACS conditions triggered lasting shifts in corticomotor excitability several minutes after the end of tACS. Only theta AM-tACS triggered a transient increase in the first minute after AC cessation. There were also no offline effects on cortical activity in the EEG measurements. It is possible that five minutes of tACS are not sufficient to induce lasting corticomotor plasticity. Although longer stimulation durations could potentially yield more persistent after-effects, this relationship is not always linear. For instance, transcranial direct current stimulation shows inconsistent gains in after-effects with extended stimulation times (Agboada et al., 2019; Samani et al., 2019), while transcranial static magnetic field stimulation produced stronger and longer-lasting reductions in motor cortex excitability with 30-minute protocols compared to shorter ones (Dileone et al., 2018).

### Phase specificity of tACS effects on corticomotor excitability

Corticospinal excitability was modestly modulated by tACS phase, with larger MEPs at the trough than the peak. This effect emerged as a consistent trend across conditions, rather than being specific to a particular frequency or waveform. Previous tACS–TMS studies that sampled multiple phases generally found maximal modulation at intermediate slopes rather than at peaks or troughs. For 20 Hz sin-tACS, Guerra et al. (2016) reported phase-dependent suppression in specific bins, with small and inconsistent peak–trough contrasts. Nakazono et al. (2016) observed facilitation around 90° during 20 Hz but not 10 Hz sin-tACS. Using closed-loop targeting, Raco et al. (2016) demonstrated phase-dependent effects peaking at intermediate phases, with substantial inter-individual variability. The same was the case in a study using cosine fits across eight bins (Schilberg et al., 2018). As our study compared only peaks and troughs, we may have underestimated phase-dependent modulation. This could account for the modest main effect of phase without robust condition-specific differences.

### Limitations

Several limitations of the present study should be acknowledged. First, the lack of a sham condition makes it difficult to entirely exclude the possibility that the concurrent application of TMS with tACS influenced the observed corticospinal excitability outcomes. Indeed, even brief TMS sessions can modulate cortical plasticity (Pellicciari et al., 2016). Second, although the order of stimulation blocks was counterbalanced, potential carryover effects cannot be entirely excluded, introducing some uncertainty regarding the isolated effects of each stimulation condition. Moreover, although participants reported no noticeable differences in skin sensations across tACS waveforms, peripheral confounds, such as cutaneous stimulation, cannot be fully ruled out. Finally, none of the protocols induced a suppression of corticospinal excitability which may indicate a propensity for the corticospinal excitability to react with increased excitability during stimulation. Since we only tested a very small selection of tACS protocols which only explored a very small proportion of the tACS parameter space and limited measurements to the resting motor state, there might well be certain frequencies at which sin-tACS and AM-tACS may suppress corticospinal excitability, for instance in the low-frequency range.

### Conclusions

Bipolar AM-tACS and sin-tACS of left M1_HAND_ with a frontocentral and a mid-parietal electrode modulate corticomotor excitability via distinct mechanisms. Beta sin-tACS likely acts through local beta entrainment, whereas theta AM-tACS appears to engage distributed network dynamics. Within-subject cross-waveform consistency in corticomotor facilitation points to trait-like variability in responsiveness, emphasizing the role of intrinsic physiological variables beyond local field strength. These findings highlight the need to disentangle the neurobiological mechanisms that mediate the actions of tACS and how these mechanisms are differentially engaged by the applied oscillatory tACS patterns. Advancing the mechanistic insights into the cortical target engagement will be critical for a principled optimization of tACS protocols.

## Conflict of interests

HRS has received honoraria as speaker from Sanofi Genzyme, Denmark and Novartis, Denmark, as consultant from Sanofi Genzyme, Denmark, Lophora, Denmark, and Lundbeck AS, Denmark, and as editor-in-chief (Neuroimage Clinical) and senior editor (NeuroImage) from Elsevier Publishers, Amsterdam, The Netherlands. He has received royalties as book editor from Springer Publishers, Stuttgart, Germany and from Gyldendal Publishers, Copenhagen, Denmark. The other authors have no conflict of interests.

## Funding

This work was supported by the Novo Nordisk Foundation Interdisciplinary Synergy Program 2014 [“Biophysically adjusted state-informed cortex stimulation (BASICS); NNF14OC0011413] to HRS.

## Acknowledgement

The authors thank Morten G Jønsson for his assistance with data collection.

## Figures & Figure captions

**Supplementary Figure 1.**
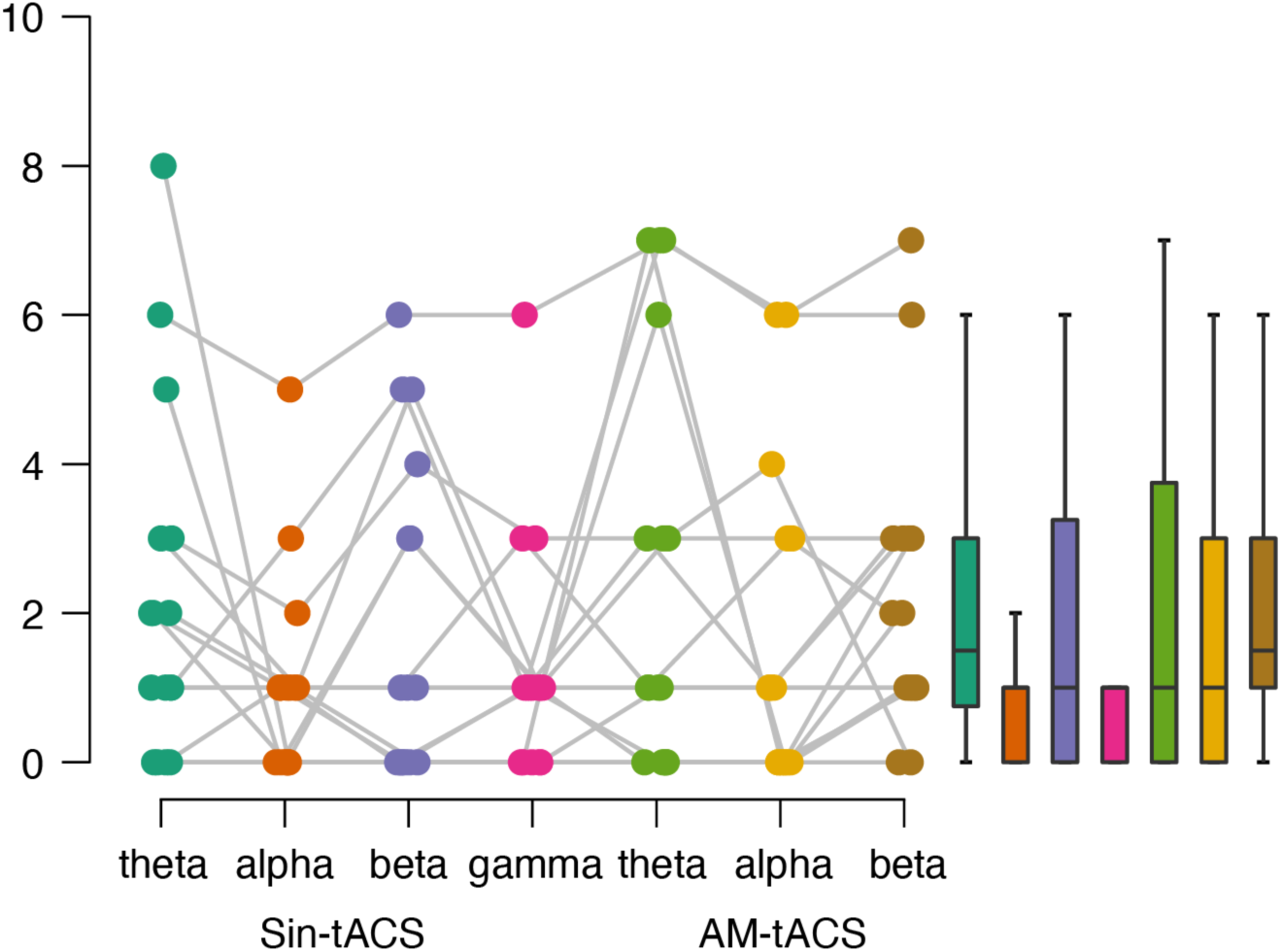
The strength and uncomfortableness of the tickling/prickling sensation by tACS in the main experiment. Each line in the left panel corresponds to individual results.

**Supplementary Figure 2.**
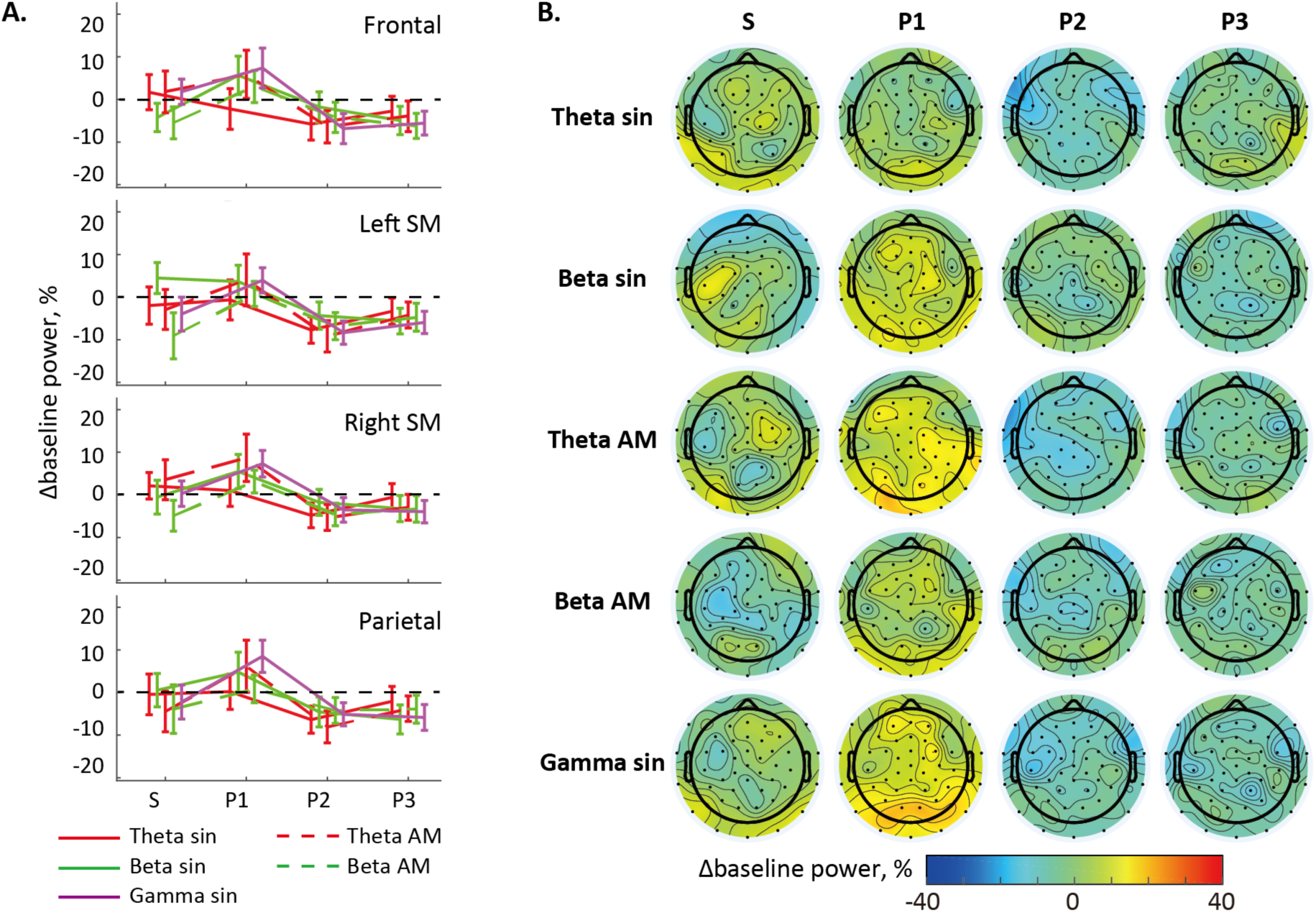
tACS-dependent changes in theta (4−6 Hz) EEG power relative to baseline. (A) Averaged EEG power within the clusters. There are no significant changes in the theta EEG power from the baseline in any time epochs. Error bars reflect standard errors of the mean. (B) Topographic representations of theta EEG power changes.

## References

[1] Agboada D, Samani MM, Jamil A, Kuo M-F, Nitsche MA. Expanding the parameter space of anodal transcranial direct current stimulation of the primary motor cortex. Sci Rep 2019;9:18185. 10.1038/s41598-019-54621-0.

[2] Akkad H, Dupont-Hadwen J, Kane E, Evans C, Barrett L, Frese A, et al. Increasing human motor skill acquisition by driving theta–gamma coupling. Elife 2021;10:e67355. 10.7554/elife.67355.

[3] Alekseichuk I, Turi Z, Amador de Lara G, Antal A, Paulus W. Spatial Working Memory in Humans Depends on Theta and High Gamma Synchronization in the Prefrontal Cortex. Curr Biol 2016;26:1513–21. 10.1016/j.cub.2016.04.035.

[4] Ali MM, Sellers KK, Fröhlich F. Transcranial alternating current stimulation modulates large-scale cortical network activity by network resonance. J Neurosci 2013;33:11262–75. 10.1523/jneurosci.5867-12.2013.

[5] Antal A, Boros K, Poreisz C, Chaieb L, Terney D, Paulus W. Comparatively weak after-effects of transcranial alternating current stimulation (tACS) on cortical excitability in humans. Brain Stimul 2008;1:97–105. 10.1016/j.brs.2007.10.001.

[6] Awiszus F. TMS and threshold hunting. Supplements to Clinical Neurophysiology 2003;56:13–23.

[7] Barzegar S, Kakies CFM, Ciupercӑ D, Wischnewski M. Transcranial alternating current stimulation for investigating complex oscillatory dynamics and interactions. Int J Psychophysiol 2025;212:112579. 10.1016/j.ijpsycho.2025.112579.

[8] Canolty RT, Edwards E, Dalal SS, Soltani M, Nagarajan SS, Kirsch HE, et al. High gamma power is phase-locked to theta oscillations in human neocortex. Science 2006;313:1626–8. 10.1126/science.1128115.

[9] Corp DT, Bereznicki HGK, Clark GM, Youssef GJ, Fried PJ, Jannati A, et al. Large-scale analysis of interindividual variability in theta-burst stimulation data: Results from the ‘Big TMS Data Collaboration.’ Brain Stimul 2020;13:1476–88. 10.1016/j.brs.2020.07.018.

[10] Daume J, Gruber T, Engel AK, Friese U. Phase-Amplitude Coupling and Long-Range Phase Synchronization Reveal Frontotemporal Interactions during Visual Working Memory. J Neurosci 2017;37:313–22. 10.1523/jneurosci.2130-16.2017.

[11] Dileone M, Mordillo-Mateos L, Oliviero A, Foffani G. Long-lasting effects of transcranial static magnetic field stimulation on motor cortex excitability. Brain Stimul 2018;11:676–88. 10.1016/j.brs.2018.02.005.

[12] Dubbioso R, Madsen KH, Thielscher A, Siebner HR. The Myelin Content of the Human Precentral Hand Knob Reflects Interindividual Differences in Manual Motor Control at the Physiological and Behavioral Level. J Neurosci 2021;41:3163–79. 10.1523/jneurosci.0390-20.2021.

[13] Feurra M, Bianco G, Santarnecchi E, Testa MD, Rossi A, Rossi S. Frequency-dependent tuning of the human motor system induced by transcranial oscillatory potentials. J Neurosci 2011;31:12165–70. 10.1523/jneurosci.0978-11.2011.

[14] Feurra M, Pasqualetti P, Bianco G, Santarnecchi E, Rossi A, Rossi S. State-dependent effects of transcranial oscillatory currents on the motor system: what you think matters. J Neurosci 2013;33:17483–9. 10.1523/jneurosci.1414-13.2013.

[15] Fricke K, Seeber AA, Thirugnanasambandam N, Paulus W, Nitsche MA, Rothwell JC. Time course of the induction of homeostatic plasticity generated by repeated transcranial direct current stimulation of the human motor cortex. J Neurophysiol 2011;105:1141–9. 10.1152/jn.00608.2009.

[16] Fröhlich F, McCormick DA. Endogenous electric fields may guide neocortical network activity. Neuron 2010;67:129–43. 10.1016/j.neuron.2010.06.005.

[17] Fröhlich F, Sellers KK, Cordle AL. Targeting the neurophysiology of cognitive systems with transcranial alternating current stimulation. Expert Rev Neurother 2015;15:145–67. 10.1586/14737175.2015.992782.

[18] Grover S, Fayzullina R, Bullard BM, Levina V, Reinhart RMG. A meta-analysis suggests that tACS improves cognition in healthy, aging, and psychiatric populations. Sci Transl Med 2023;15:eabo2044. 10.1126/scitranslmed.abo2044.

[19] Guerra A, Pogosyan A, Nowak M, Tan H, Ferreri F, Lazzaro VD, et al. Phase Dependency of the Human Primary Motor Cortex and Cholinergic Inhibition Cancelation During Beta tACS. Cereb Cortex 2016;26. 10.1093/cercor/bhw245.

[20] Helfrich RF, Schneider TR, Rach S, Trautmann-Lengsfeld SA, Engel AK, Herrmann CS. Entrainment of brain oscillations by transcranial alternating current stimulation. Curr Biol 2014;24:333–9. 10.1016/j.cub.2013.12.041.

[21] Helfrich RF, Knight RT. Oscillatory Dynamics of Prefrontal Cognitive Control. Trends Cogn Sci 2016;20:916–30. 10.1016/j.tics.2016.09.007.

[22] Herrmann CS, Rach S, Neuling T, Strüber D. Transcranial alternating current stimulation: a review of the underlying mechanisms and modulation of cognitive processes. Front Hum Neurosci 2013;7:279. 10.3389/fnhum.2013.00279.

[23] Hsu C, Liu T, Lee D, Yeh D, Chen Y, Liang W, et al. Amplitude modulating frequency overrides carrier frequency in tACS-induced phosphene percept. Hum Brain Mapp 2022;44:914–26. 10.1002/hbm.26111.

[24] Johnson L, Alekseichuk I, Krieg J, Doyle A, Yu Y, Vitek J, et al. Dose-dependent effects of transcranial alternating current stimulation on spike timing in awake nonhuman primates. Sci Adv 2020;6:eaaz2747. 10.1126/sciadv.aaz2747.

[25] Kasten FH, Duecker K, Maack MC, Meiser A, Herrmann CS. Integrating electric field modeling and neuroimaging to explain inter-individual variability of tACS effects. Nat Commun 2019;10:27-Nov. 10.1038/s41467-019-13417-6.

[26] Khanna P, Carmena JM. Neural oscillations: beta band activity across motor networks. Curr Opin Neurobiol 2015;32:60–7. 10.1016/j.conb.2014.11.010.

[27] Krause MR, Vieira PG, Csorba BA, Pilly PK, Pack CC. Transcranial alternating current stimulation entrains single-neuron activity in the primate brain. Proc Natl Acad Sci USA 2019;116:5747–55. 10.1073/pnas.1815958116.

[28] Li L, Gainey MA, Goldbeck JE, Feldman DE. Rapid homeostasis by disinhibition during whisker map plasticity. Proc Natl Acad Sci 2014;111:1616–21. 10.1073/pnas.1312455111.

[29] Lisman JE, Jensen O. The Theta-Gamma Neural Code. Neuron 2013;77:1002–16. 10.1016/j.neuron.2013.03.007.

[30] Miniussi C, Bortoletto M. Harnessing neural variability: Implications for brain research and non-invasive brain stimulation. Neurosci Biobehav Rev 2025;176:106312. 10.1016/j.neubiorev.2025.106312.

[31] Misonou H. Homeostatic Regulation of Neuronal Excitability by K+ Channels in Normal and Diseased Brains. Neurosci 2010;16:51–64. 10.1177/1073858409341085.

[32] Moliadze V, Antal A, Paulus W. Boosting brain excitability by transcranial high frequency stimulation in the ripple range. J Physiol 2010;588:4891–904. 10.1113/jphysiol.2010.196998.

[33] Mölle M, Born J. Slow oscillations orchestrating fast oscillations and memory consolidation. Prog Brain Res 2011;193:93–110. 10.1016/b978-0-444-53839-0.00007-7.

[34] Nakazono H, Ogata K, Kuroda T, Tobimatsu S. Phase and Frequency-Dependent Effects of Transcranial Alternating Current Stimulation on Motor Cortical Excitability. Plos One 2016;11:e0162521. 10.1371/journal.pone.0162521.

[35] Negahbani E, Kasten FH, Herrmann CS, Fröhlich F. Targeting alpha-band oscillations in a cortical model with amplitude-modulated high-frequency transcranial electric stimulation. Neuroimage 2018;173:12-Mar. 10.1016/j.neuroimage.2018.02.005.

[36] Noury N, Siegel M. Phase properties of transcranial electrical stimulation artifacts in electrophysiological recordings. Neuroimage 2017;158:406–16. 10.1016/j.neuroimage.2017.07.010.

[37] Noury N, Hipp JF, Siegel M. Physiological processes non-linearly affect electrophysiological recordings during transcranial electric stimulation. Neuroimage 2016. 10.1016/j.neuroimage.2016.03.065.

[38] Oldfield RC. The assessment and analysis of handedness: The Edinburgh inventory. Neuropsychologia 1971;9:97–113. 10.1016/0028-3932(71)90067-4.

[39] Pellicciari MC, Miniussi C, Ferrari C, Koch G, Bortoletto M. Ongoing cumulative effects of single TMS pulses on corticospinal excitability: An intra- and inter-block investigation. Clin Neurophysiol 2016;127:621–8. 10.1016/j.clinph.2015.03.002.

[40] Pozdniakov I, Vorobiova AN, Galli G, Rossi S, Feurra M. Online and offline effects of transcranial alternating current stimulation of the primary motor cortex. Sci Rep-Uk 2021;11:3854. 10.1038/s41598-021-83449-w.

[41] Raco V, Bauer R, Tharsan S, Gharabaghi A. Combining TMS and tACS for Closed-Loop Phase-Dependent Modulation of Corticospinal Excitability: A Feasibility Study. Front Cell Neurosci 2016;10:143. 10.3389/fncel.2016.00143.

[42] Ridding MC, Ziemann U. Determinants of the induction of cortical plasticity by non-invasive brain stimulation in healthy subjects. J Physiol 2010;588:2291–304. 10.1113/jphysiol.2010.190314.

[43] Riddle J, McFerren A, Frohlich F. Causal role of cross-frequency coupling in distinct components of cognitive control. Prog Neurobiol 2021;202:102033. 10.1016/j.pneurobio.2021.102033.

[44] Riddle J, Frohlich F. Targeting neural oscillations with transcranial alternating current stimulation. Brain Res 2021;1765:147491. 10.1016/j.brainres.2021.147491.

[45] Romer SH, Deardorff AS, Fyffe REW. Activity-dependent redistribution of Kv2.1 ion channels on rat spinal motoneurons. Physiol Rep 2016;4:e13039. 10.14814/phy2.13039.

[46] Samani MM, Agboada D, Jamil A, Kuo M-F, Nitsche MA. Titrating the neuroplastic effects of cathodal transcranial direct current stimulation (tDCS) over the primary motor cortex. Cortex 2019;119:350–61. 10.1016/j.cortex.2019.04.016.

[47] Sauseng P, Peylo C, Biel AL, Friedrich EVC, Romberg-Taylor C. Does cross-frequency phase coupling of oscillatory brain activity contribute to a better understanding of visual working memory? Br J Psychol 2019;110:245–55. 10.1111/bjop.12340.

[48] Schilberg L, Engelen T, Oever ST, Schuhmann T, Gelder B de, Graaf TA de, et al. Phase of beta-frequency tACS over primary motor cortex modulates corticospinal excitability. Cortex 2018;103:142–52. 10.1016/j.cortex.2018.03.001.

[49] Schutter DJLG, Hortensius R. Brain oscillations and frequency-dependent modulation of cortical excitability. Brain Stimul 2011;4:97–103. 10.1016/j.brs.2010.07.002.

[50] Solomon EA, Kragel JE, Sperling MR, Sharan A, Worrell G, Kucewicz M, et al. Widespread theta synchrony and high-frequency desynchronization underlies enhanced cognition. Nat Commun 2017;8:1704. 10.1038/s41467-017-01763-2.

[51] Thielscher A, Antunes A, Saturnino GB. Field modeling for transcranial magnetic stimulation: A useful tool to understand the physiological effects of TMS? vol. 2015, 2015, p. 222–5. 10.1109/embc.2015.7318340.

[52] Wischnewski M, Schutter DJLG, Nitsche MA. Effects of beta-tACS on corticospinal excitability: A meta-analysis. Brain Stimul 2019;12:1381–9. 10.1016/j.brs.2019.07.023.

[53] Wischnewski M, Alekseichuk I, Opitz A. Neurocognitive, physiological, and biophysical effects of transcranial alternating current stimulation. Trends Cogn Sci 2023;27:189–205. 10.1016/j.tics.2022.11.013.

[54] Witkowski M, Cossio EG, Chander BS, Braun C, Birbaumer N, Robinson SE, et al. Mapping entrained brain oscillations during transcranial alternating current stimulation (tACS). Neuroimage 2015. 10.1016/j.neuroimage.2015.10.024.

[55] Yanagisawa T, Yamashita O, Hirata M, Kishima H, Saitoh Y, Goto T, et al. Regulation of motor representation by phase-amplitude coupling in the sensorimotor cortex. J Neurosci 2012;32:15467–75. 10.1523/jneurosci.2929-12.2012.

[56] Ziemann U, Siebner HR. Inter-subject and Inter-session Variability of Plasticity Induction by Non-invasive Brain Stimulation: Boon or Bane? Brain Stimul 2015;8:662–3. 10.1016/j.brs.2015.01.409.

